# Integrated Analysis of Cell Shape and Movement in Moving Frame

**DOI:** 10.1101/2020.11.04.369033

**Authors:** Yusri Dwi Heryanto, Chin-Yi Cheng, Yutaka Uchida, Kazushi Mimura, Masaru Ishii, Ryo Yamada

## Abstract

The cell movement and morphological change are two interrelated cellular processes. The integrated analysis is needed to explore the relationship between them. However, it has been challenging to investigate them as a whole. The cell trajectory can be described by its speed, curvature, and torsion. On the other hand, the 3-Dimensional (3D) cell shape can be studied by using a shape descriptor such as Spherical harmonic (SH) descriptor which is an extension of Fourier transform in 3D space.

To integrate these shape-movement data, we propose a method to use a parallel-transport (PT) moving frame as the 3D-shape coordinate system. This moving frame is purely determined by the velocity vector. On this moving frame, the movement change will influence the coordinate system for shape analysis. By analyzing the change of the SH coefficients over time in the moving frame, we can observe the relationship between shape and movement.

We illustrate the application of our approach using simulational and real datasets in this paper.

## INTRODUCTION

The cell movement and morphological change are highly integrated. They share many biological mechanisms controlled by the cytoskeleton, cell membrane, membrane proteins, and extracellular matrix(Friedl and Wolf, 2009; Ridley, 2003). Almost all forms of the cell active movement need forces that are generated from dynamic shape change(Lämmermann and Sixt, 2009). Moreover, the difference in the shapes and sizes of motile cells reflects their movement pattern(Keren et al., 2008; Lämmermann and Sixt, 2009). To study the shape-movement of the cell as a whole connected process, the integrated analysis is essential.

The first step in the shape analysis is shape normalization. Cells are 3D objects that usually have arbitrary spatial positions, directions, and scales in the 3D-space. However, these shapes may be variations of the same shape and should be recognized as the same one. Shape normalization will put the object in the common frame of reference and make shape analysis more robust to these isometric transformations (i.e. translation, scale, rotation, and reflection).

From all isometric transformations, orientation is usually the hardest one to be normalized. Some shapes with a distinguishable axis of orientation such as an ellipsoid are easy to orient using their longest axis as the axis of orientation. Yet, many shapes do not have a clear axis of orientation. Many studies investigated this topic with different approaches. The most well-known approach is the principal component analysis based approach(Shilane et al., 2004). However, these methods do not provide robust normalization and can produce poor alignments(Kazhdan et al., 2003; Kazhdan, 2007). Another approach is to transform each shape into a function and then calculate the rotation that minimizes the distance between the two functions(Makadia and Daniilidis; Makadia et al., 2004). The weakness of this approach is the alignment results are inconsistent and depend on the initial pose of the model.

In this paper, we propose a novel method to solve this alignment problem and integrate the analysis of shape and movement at the same time by aligning the 3D objects using the parallel-transport (PT) moving frame(Bishop, 1975; Hanson and Ma, 1995). This moving frame is constructed primarily using the velocity vector. Therefore, the PT moving frame is more natural for moving biological objects as the frame of reference. For illustration, we can orient a car or a plane using an axis that is parallel with their movement direction. In other words, the movement direction become the axis of the basis. If there is some movement change, it will also change the basis of the shape. Hence, this approach can bridge the analysis of shape and movement. To illustrate our approach, we use both of simulated and real datasets.

### Related work

The study of dynamic morphological change of motile cells was started from the study of 2-dimensional (2D) shapes. The 2D migration is characterized by the unilateral adhesion to a substrate and a spread-out cell shape with a leading lamellipod(Ridley, 2003; Keren et al., 2008). Lee et al. proposed the graded radial extension (GRE) model which described the locomotion and shape change of keratocytes that consistent with the half-moon shape and its gliding motion (Lee et al., 1993). This model assumed that the local cell extension (protrusion or retraction) are perpendicular to the cell edge and the cell maintains its shape and size during moving by a graded distribution (e.g. a maximum near the cell midline to a minimum towards the sides). Further study found that keratocytes inhabit a low dimensional shape space(Keren et al., 2008). The Cellular Potts Model (CPM) is another popular model for dynamic, irregular, and highly fluctuating cell shapes(Graner and Glazier, 1992; Niculescu et al., 2015; Rens and Edelstein-Keshet, 2019). A cell in CPM is defined over a multiple lattice sites region with the constraint on the cell area. In this model, the cell is exposed to an energy function from adhesion between cells and resistance of cells to volume changes. Then, the cell motion and shape change comes from the minimization of this energy function(Marée et al.).

Even though the study on 2D space is important, the cells live and move in the 3D environments. A 3D shape is more challenging to study due to higher degrees of freedom. A common approach is to represent the 3D object surface as a shape descriptor. By examining the change of the shape descriptors over time, we can study the dynamic of the shape change.

Spherical harmonic (SH) descriptor is a widely used descriptor to study the 3D cell shape(Shen et al., 2009; Ducroz et al., 2012; Du et al., 2013; Medyukhina et al., 2020). It is a spherical analogue of the 1D Fourier series. It considers a surface as a function on the unit sphere which can be represented as a set of unique coefficients. In the contexts of shape-movement analysis, SH was used to analyzed amoeboid cell motion(Ducroz et al., 2012; Du et al., 2013) and to perform shape classification of motile cells(Medyukhina et al., 2020). As mentioned before, any shape analysis including the SH descriptor needs to be robust to rotational transformation. A rotation invariant version of the SH descriptor is commonly used to address this problem(Kazhdan et al., 2003). However, this method suffers information lost because they described a shape in a transformation invariant manner (e.g. disregard the effect of rotation)(Kazhdan et al, 2003).

In this paper, we propose a shape-movement analysis using the SH descriptor in the parallel-transport moving frame. In contrast to the previous approach, we keep the effect of rotation by aligned all the shapes in the common frame of reference based on the parallel-transport moving frame. Another contribution from our approach, we can observe and quantify the relationship between the movement and the shape of the cell. It is possible because the moving frame is constructed from the velocity vector of the object.

## METHODS

### Overview

The schematic diagram of our approach is shown in FIG. 1. The trajectory of the cell was smoothed using Gaussian Process regression. From this smooth curve, we constructed moving frames at each sample point and extracted the trajectory properties such as the speed, curvature, and torsion at each sample point. For each 3D object from each timepoint, we change the coordinate basis from the standard basis to the moving frame basis. In this moving frame, the cell direction become the new x-axis. We performed the SH decomposition on this new basis to obtain SH coefficients. By analyzing the change of these coefficients over time, we can observe the dynamic of shape change when it was moving. We performed the statistical analysis of the shape and movement features to study their relationship.

**Fig. 1.**
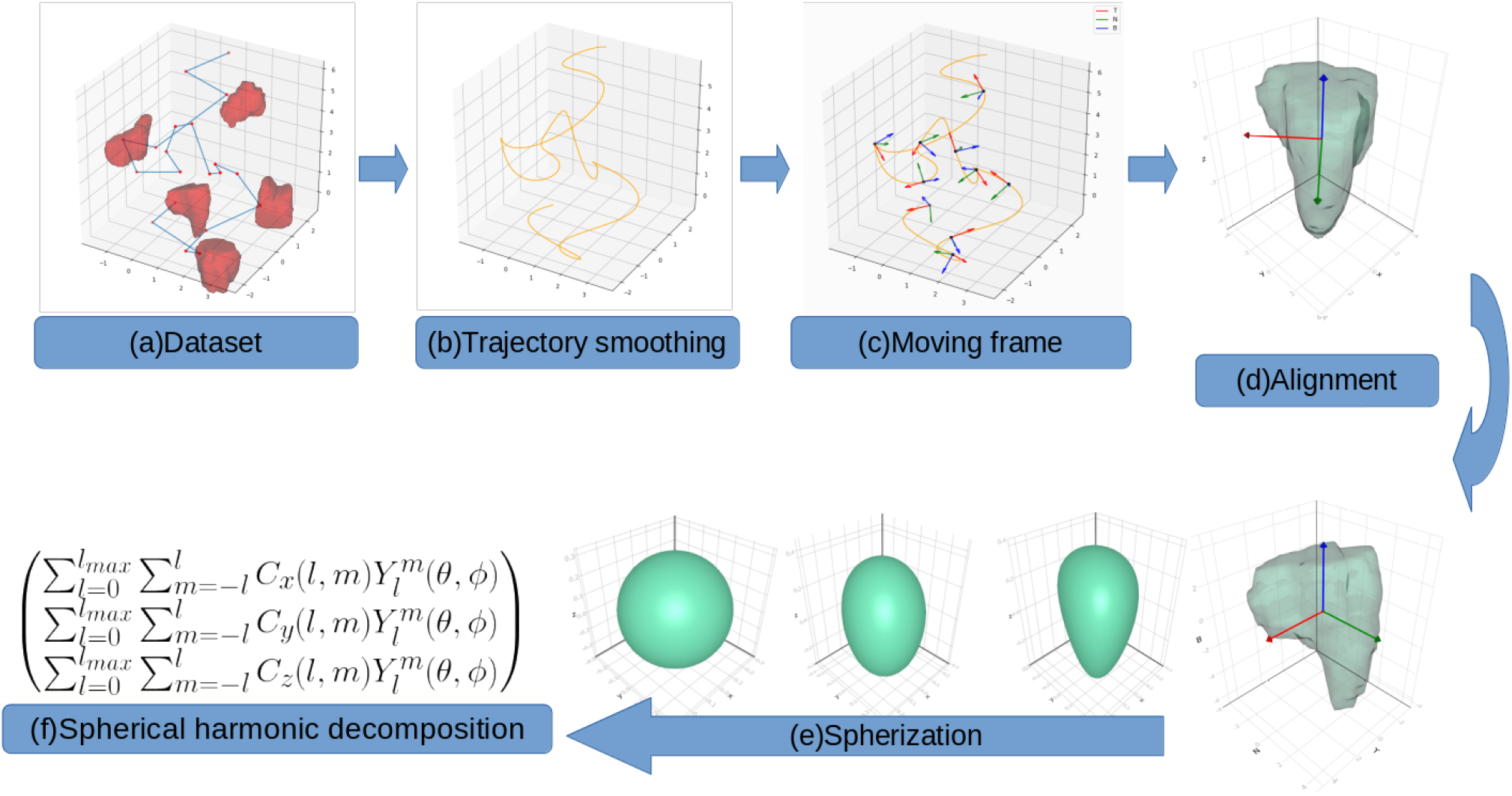
The schematic diagram of integrated analysis of shape and movement in moving frame.

### Trajectory and Moving Frame of the Mass Center

#### Gaussian Process smoothing

We obtained the trajectory by calculating the mass center of the 3D object at every timepoint. The mass center of the object is defined as the arithmetic mean of their vertices. Thus, we had (*x*(*t*), *y*(*t*), *z*(*t*)) as the position of the mass center at the time *t*. Then, we defined *s_x_* = (*x*(1), .., *x*(*n*)), *s_y_* = (*y*(1), .., *y*(*n*)), and *s_z_* = (*z*(1), .., *z*(*n*)). For each sequence *s_i_*; *i* = *x, y, z*, we standardized the sequence so that it has mean zero and standard deviation one. From hereafter, the subscript *i* refers to one of the element of *x, y, z*, unless otherwise stated.

We modeled each of the sequence as 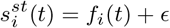 where 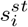 is a standardized sequence. The function *f_i_*(*t*) is an underlying smooth function and ∊ is a Gaussian noise with mean zero and variance *σ*^2^.

The function *f_i_*(*t*) can be approximated by Gaussian Process (GP) Regression(Rasmussen and Williams, 2006). In the GP regression, we have mean function and kernel function as hyperparameters. As our sequence had zero mean, we chose the zero mean function. And because we need a smooth function, we used Squared-Exponential (SE) kernel function.

The SE kernel itself has two parameters: (1) the lengthscale parameter that determines how far the influence of one sample point to its neighbor points, (2) the variance parameter that determines the mean distance from the function’s mean.

We used the function *f_i_*(*t*) to interpolate new data points between two consecutive elements of sequence 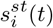 and 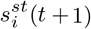. The number of new data points determines the smoothness of the new sequence.

Next, we performed the inverse transform of standardization by multiply 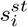 by the standard deviation of *s_i_*, then add it with the mean of *s_i_* to produce a near-smooth sequence *r_i_*. Then, we defined **r**(*t*) = (*r_x_*(*t*), *r_y_*(*t*), *r_z_*(*t*)) as the smooth trajectory.

#### Parallel transport moving frame

One of the main ideas in our approach is a frame of reference which is moving along with the curve and telling us the main directions of the movement (Fig S1). We made this frame of reference using parallel-transport (PT) moving frame algorithm from Hanson and Ma (1995). In brief, we calculated a tangent vector **T_i_** for each point *i* on the curve. Then, set an initial normal vector **N_0_** which is perpendicular to the first tangent vector **T_0_**. For each sampled point, we calculate the cross product of **U** = **T_i_** × **T_i+1_**. If the length ||U|| = 0 (i.e. both vectors are parallel) then **N_i+1_** = **N_i_**. Otherwise, to obtain **N_i+1_**, we rotate **N_i_** around the vector **U** by the angle *θ* = **T_i_** · **T_i+1_** using the rotation matrix *Rot*(**U**, *θ*) defined on Hanson and Ma (1995). The complete algorithm is shown in Algorithm 1.

#### Orientation normalization

We change the coordinate basis from the standard basis to the PT moving frame basis using simple linear algebra transformation:

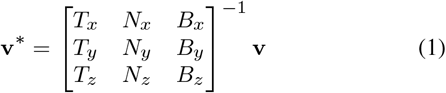

where **v**\* and **v** are the coordinates of the vertex in moving frame and standard basis respectively. The subscript *x, y*, and *z* here are the first, second, and third component of the vectors **T**, **N**, and **B**.

#### Speed, curvature, and torsion

From the smooth trajectory, we can extract the trajectory features such as speed, curvature, and torsion as defined on Patrikalakis and Maekawa (2009). In summary, speed is the distance traveled per unit of time. Curvature measures the failure of a trajectory to be a straight line, while torsion measures the failure of a trajectory to be planar. Taken together, the curvature and the torsion of a space curve are analogous to the curvature of a plane curve.

##### Algorithm 1

Parallel transport moving frame algorithms

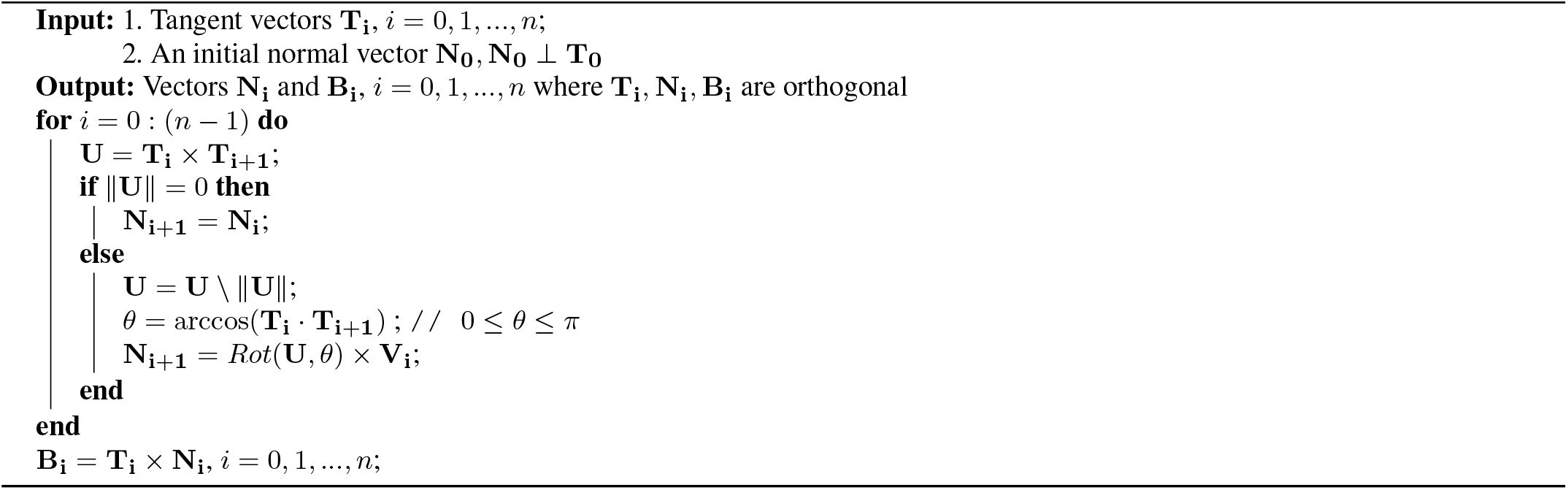

### Spherical Harmonic Decomposition

After orientation normalization, shapes were decomposed Spherical Harmonic (SH) transform. To perform SH decomposition, we need to map the object surface to the unit sphere. Before it, we normalize the volume of all objects to one.

#### Spherical parameterization

We used the mean-curvature flow spherical parameterization method from Kazhdan(Kazhdan et al., 2012) to maps the object surface to a unit sphere. The result of spherical parameterization is a continuous and uniform mapping between a point on the object surface and a pair of the latitudinal-longitudinal coordinate (*θ, ϕ*) on a unit sphere:

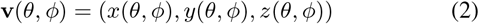

where (*x*(*θ, ϕ*), *y*(*θ, ϕ*), *z*(*θ, ϕ*)) is the Cartesian vertex coordinates.

#### Spherical Harmonic expansion

On the unit sphere, each of the (*x*(*θ, ϕ*), *y*(*θ, ϕ*), *z*(*θ, ϕ*) can be approximated using the real form spherical harmonic(SH) series:

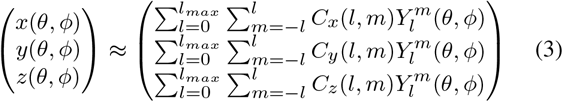

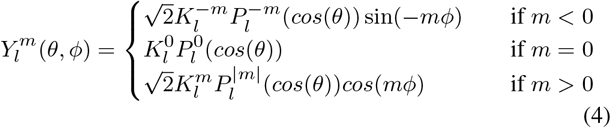

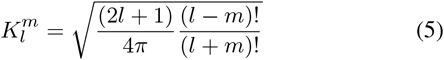

where 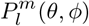 is the associated Legendre polynomial.

The coefficient of degree l, order m, *C*(*l, m*) can be obtained using standard least-square estimation. Using *x*(*θ, ϕ*) as an example, assume that the number of vertices is *n* and *x_i_* = *x*(*θ_i_, ϕ_i_*). We need to find the coefficients *C_x_* = (*c*_1_, *c*_2_,⋯, *c_k_*)^*T*^ where *c_j_* = *C_x_*(*l, m*). The index *j* for *l* ∈ (0,⋯, *l_max_*), *m* ∈ (−*l*,⋯, *l*) is obtained from the equation *j* = *l*^2^ + *l* + *m* + 1. We can obtain the coefficients by solving equation 6.

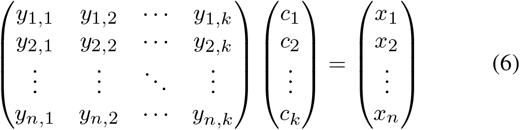

where 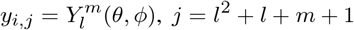, and *k* = *l_max_* + 1. After obtaining the coefficients for each of *C_x_, C_y_*, and *C_z_*, we can bundle it into one feature vector of the shape, *C* = (*C_x_, C_y_, C_z_*).

#### Shape characteristics measures

Each coefficient of the expansion retains a shape information corresponding to a particular spatial frequency. The increasing degree of *l* describes the finer scales of shape information. The direction of shape changes can be detected in each of three sets of coefficients, especially *C_x_*(*l* = 1, *m* = 1) for deformation in the x-direction, *C_y_*(1,0) for y-direction, and *C_z_* (1, − 1) for z-direction. If *C_x_*(1, 1) = *C_y_*(1, *m* = 0) = *C_z_*(1, −1) and the other coefficients are zero then the object is a perfect sphere. Based on these three coefficients, we defined three eccentricity index:

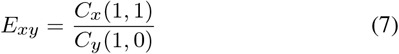

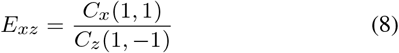

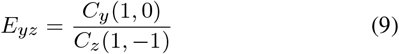

*E_xy_*, *E_xz_*, and *E_yz_* are the measurements of how much the object deviates from being sphere in xy, xz, and yz plane, respectively (Fig. S2). We can exclude the *E_yz_* because the calculation of *E_xy_* and *E_xz_* already contains all of the eccentricity information.

The difference between shape *i* and *j*, *d*(*i, j*), can be calculated using any distance metrics intended for real-valued vector spaces. The most common is the *L*_2_ norm distance:

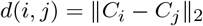

where *C_i_* and *C_j_* are SH coefficients for shape *i* and *j*, respectively. Here, we calculated the the rate of shape change which is defined as *d*(*t, t* + 1) where *t, t* + 1 are two consecutive timepoints.

#### Statistical analysis

For each cell in the real dataset, we calculated the median, median absolute deviation, 25th percentile, and 75th percentile of the speed, curvature, torsion, shape change rate, *E_xy_*, and *E_xz_* from all timepoints. We also counted the number of peaks from the curvature and torsion graph. These values are used as features (26 features in total). Then, we standardized all these features to have mean zero and standard deviation one. To show the correlation from each features, we calculated Kendall rank correlation coefficient.

We use UMAP (McInnes et al., 2018) for visualization in a 2D plot. The UMAP parameters that we used: number of neighbors = 20, number of components = 2, minimum distance = 0.1, and the Euclidean distance as metric.

K-nearest neighbor (KNN) classifier was trained on the shape-movement features on the PT moving frame and on the standard basis to show the performance of our approach. We varied the hyper-parameter *k* = 3, 5, 10, 15, 20, 25. A stratified 10-fold crossvalidation was used to give a set of 10 accuracy scores for each hyper-parameter *k*. From each iteration of cross-validation, we calculated the difference of accuracy scores between these two approaches. The One sample T-test was performed to test whether the accuracy difference was significantly different from zero or not.

### Datasets

#### Simulational dataset

We manually created a 3D cell object that moves along a path using open-source software Blender (Community, 2018). The dataset consists of 250 time points and a 3D object at each time point. The 3D objects were saved as triangular mesh objects. In our simulation, the cell moves in the accelerate-decelerate-accelerate-decelerate pattern. The cell starts from a near-spherical shape, protruding a pseudopod when accelerating, and back to near-spherical shape when decelerating (Movie 1,Movie 2a). The volume of the cell is fixed to be one. We chose the shape object at timepoint *t* = {1, 10, 20,…, 250} as the observation data. The rest of the data were used as the holdout data. The true trajectory of the cell is defined as the center of the mass of the cell at each of 250 timepoints.

#### Real dataset

The real dataset consists of 3D objects from microscope images of neutrophils. We isolated the neutrophils from the bone marrow of LysM-EGFP mice. Erythrocytes were excluded from harvested bone marrow cells using ACK lysing buffer and density gradient centrifugation(800xG for 20 minutes) using 62.5 percent percoll. The isolated neutrophils were mixed with collagen with 3 × 10^4^ cells/μl. Next, we dropped collagen-cell mixture(10 ul) on the glass bottom dish. After the collagen-cell mixture turned into a gel, we add culture medium to the dish. The dish was incubated at 37 °C for 2 hours. Before the imaging, the culture medium was replaced with the imaging medium. After the cells were stimulated with Granulocyte-Macrophage Colony-Stimulating-Factor(GM-CSF) 25 ng ml^−1^, Lipopolysaccharide(LPS) 10 μg ml^−1^, or (Phorbol 12-myristate-13-acetate(PMA) 1 μgml^−1^, we immediately performed imaging for 90 minutes at 1-minute intervals using two-photon excitation microscope (Nikon A1R MP). The imaging was performed at 45 μm (15 stacks) in 3 μm steps in the Z-axis direction.

We analyzed the cells that were captured in a minimal 6 consecutive timepoints(i.e. 233, 398, 293, and 166 cells in saline, GM-CSF, LPS, and PMA group, respectively).

### Method Implementation

The above method was developed in Julia Programming Language(Bezanson et al., 2017). We used Gaussian Processes.jl(Fairbrother et al., 2018) package for the Gaussian Process smoothing. For SH coefficient calculation, we used Julia wrapper for the GNU Scientific Library (GSL)(Galassi, 2009).

For the SE kernel parameter, we set the lengthscale=*e*^1.0^ and variance = 1.0. We note that the choice of the parameters is not necessarily optimal, but it gives good modeling results in our simulation, as will be shown later. For the SH decomposition, we used *l_max_* = 6.

## RESULTS

### Simulational dataset

The smooth trajectory from the observation data is shown in FIG. 2a. It is difficult to see any difference due to almost perfect reconstruction (The mean squared error = 1.203). The quality of the smooth trajectory degrades around t=0 and t=250. This is probably due to the Gaussian Processes had fewer data to process around the boundary (i.e. no data at *t* < 0 or *t* > 250).

**Fig. 2.**
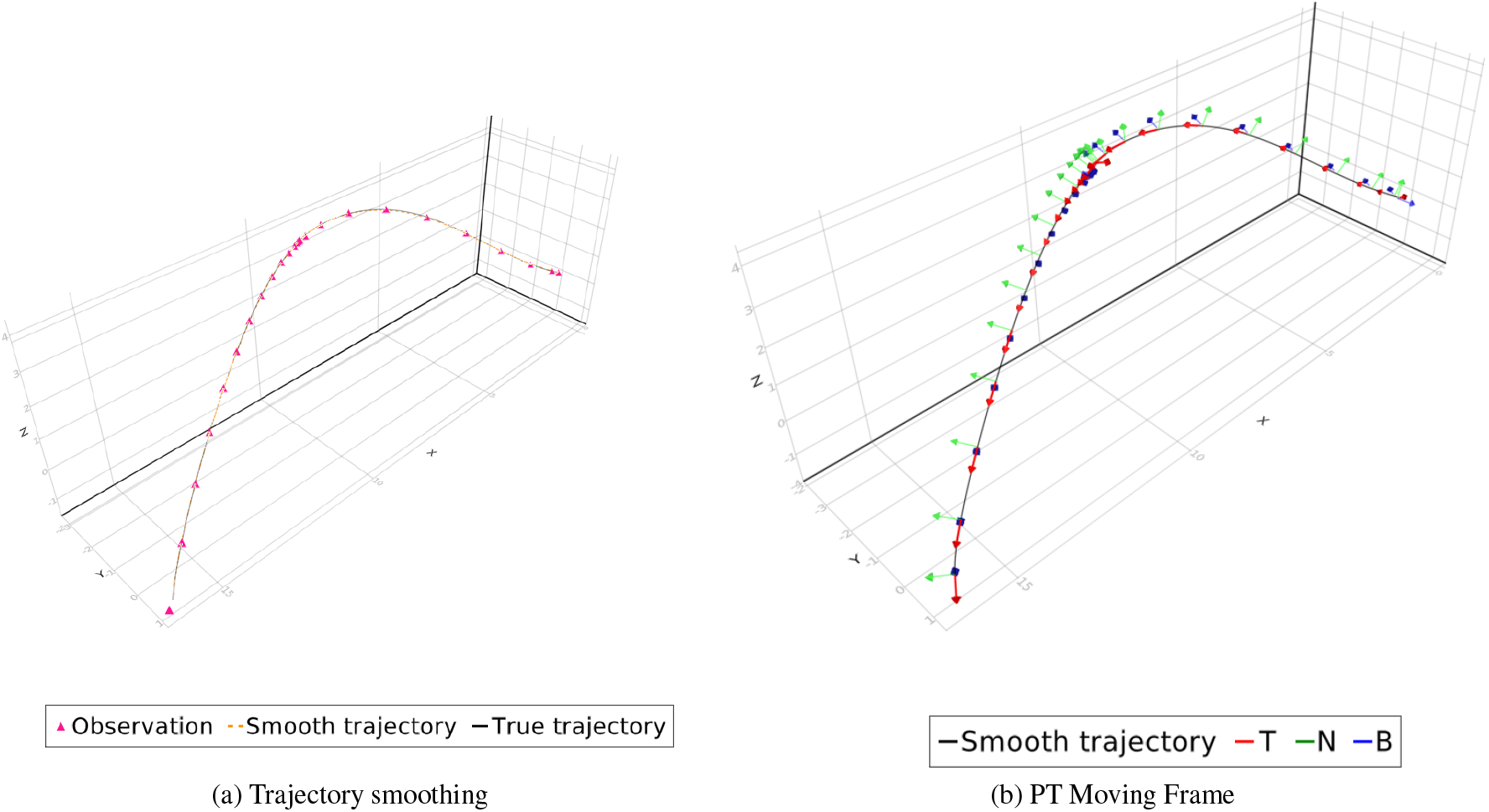
Trajectory of simulated cell.(a)The Gaussian Process reconstructed the original trajectory from the observation.(b) The vectors **T** together with the vectors **N** and **B** construct moving frames for each point on the curve.

We constructed the PT moving frames on this smooth trajectory (FIG. 2b). Orientation normalization was performed using PT frames as the canonical frame of reference (FIG. 3). After reorientation, we observed that the cell protrude its pseudopod in the direction of the cells (Movie 2b).

**Fig. 3.**
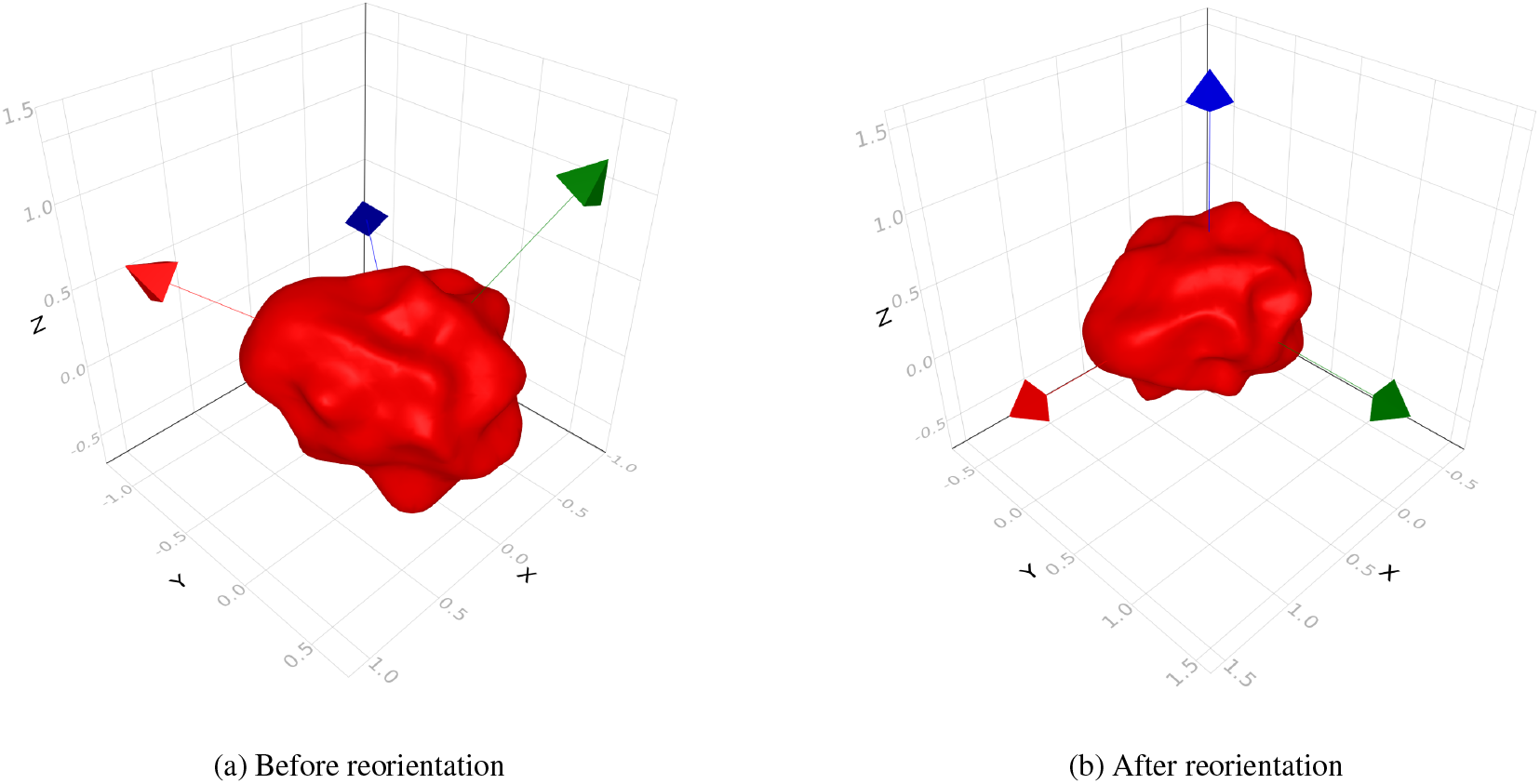
Reorientation of shape using PT moving frame

The shape-movement features of the simulational cell is shown on FIG. 4. In these plots, the features of the original data are also shown. The pattern that we obtained from observation data are similar to the original data, even though the magnitude of the peak is different. These differences due to the data from observational data is smaller than the original data.

**Fig. 4.**
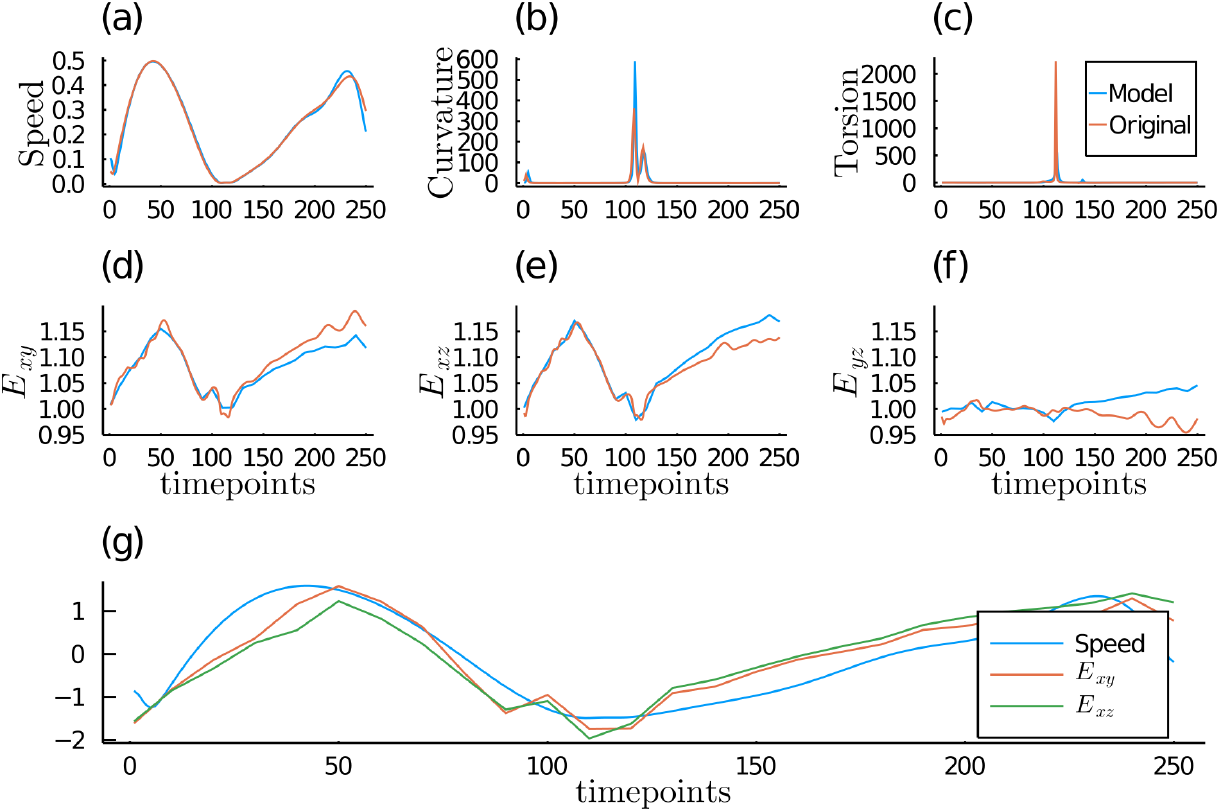
The shape and movement pattern of the cell over time from observation data and its comparison with the original data. (a,b,c) The movement features of the cell are characterized by its speed, curvature, and torsion. (d,e,f) The global shape features of the cell is represented by the ratio of spherical harmonic coefficients. (g) The relation between shape and movement of the cell can be seen in this graph. Here, we standardized the graph of speed, *E_xy_*, and *E_xz_* for easy comparison.

On the upper part of FIG. 4, the movement behavior of the cell is shown on the speed, curvature, and torsion graph. The bimodal graph of the speed graph indicates the accelerate-decelerate-accelerate-decelerate pattern. Meanwhile, the peak on the curvature and torsion graph indicate the change of the direction between *t* = 100 and *t* = 130.

On the middle part of FIG. 4, the global deformation patterns of the cell are shown as the eccentricity of the shape. Here, *E_xy_* and *E_xz_* changed as time progressed and had values more than one in the majority of the time. In contrast, the value *E_yz_* did not vary a lot from one. These finding indicate that the cell had ellipsoid shape with the longer axis on the direction of the movement. Furthermore, the relation between shape and movement can be seen on lower part of the FIG. 4. In this plot, the speed, the *E_xy_*, and *E_xz_* are standardized for comparison purpose. We can easily identify the similarity of the bimodal pattern on the speed, the *E_xy_*, and *E_xz_* plot.

### Real data

The UMAP was utilized to visualize the shape-movement features of the real dataset on 2-dimensional plot. The FIG. 5a and the FIG. 5b shows the 2D plot of the features from PT moving frame and the standard basis. Qualitatively, the features obtained on the PT moving frame separate each group better than the features from the standard basis.

**Fig. 5.**
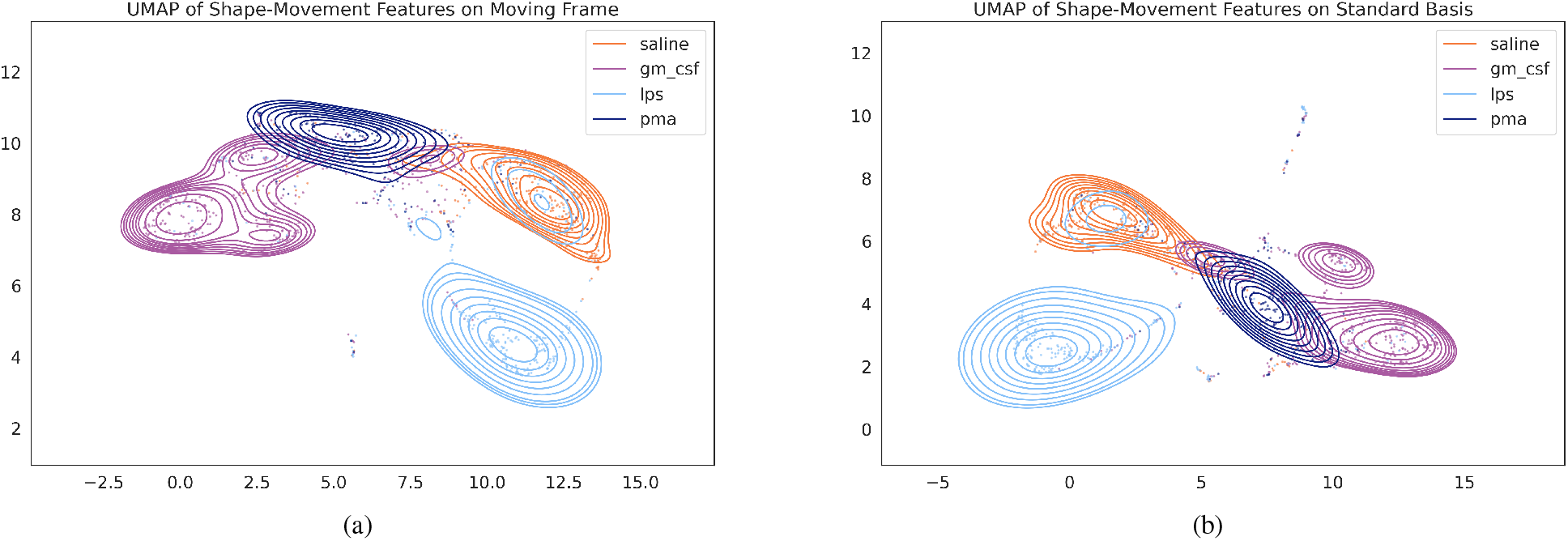
The Shape-Movement features embedding on 2D space using UMAP. We utilized UMAP to visualize the high-dimensional shape-movement features on (a) PT moving frame and on (b) Standard basis.

Quantitatively, we performed KNN classifier on the features from the PT moving frame and the standard basis. The FIG. 6a show the mean and the standard deviation of the KNN classifier accuracy. The majority of the accuracy scores on the PT moving frame are superior to the standard basis. Using One Sample T-test, the accuracy difference is significant (i.e. p-value < 0.05) on *k* = 15, 20, 25 (FIG. 6b).

**Fig. 6.**
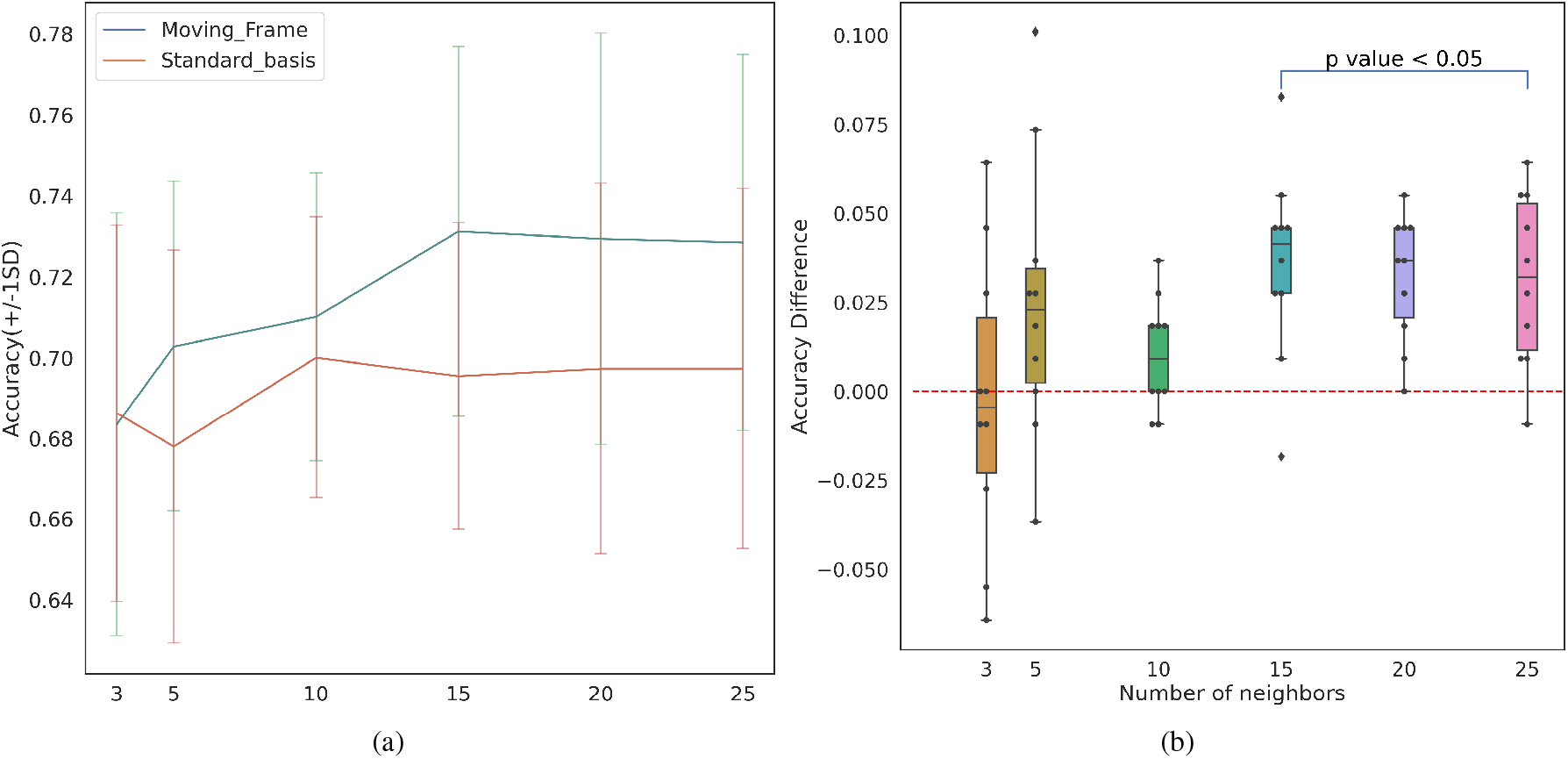
The analysis of shape-movement features on PT moving frame. (a) The comparison of accuracy of KNN classifier across the 10-folds cross-validation trained on PT moving frame and on standard basis. (b) The accuracy difference of KNN classifier on PT moving frame and on standard basis. The one sample t-test showed that the difference was significantly different from zero for *k* = 15, 20, 25

On FIG. 7, we plot the importance of each feature on the accuracy of KNN classifier. The most importance features come from the speed features which is the number of the peak on curvature graph, the number of the peak on torsion graph, the 75th percentile and median absolute deviation of the speed. Here, the shape features play a minor role to classify the sample. We can see the relationship between each feature from the Fig 8. We found some interesting observations from it. Torsion and curvature features are negatively correlated with the speed features. It suggests that the cells that are moving faster are inclined to move straight(i.e. rarely change their direction and low magnitude of direction change). We also found that the fast cells had the rate of shape change higher and protrude more along its movement direction than the slower cells.

**Fig. 7.**
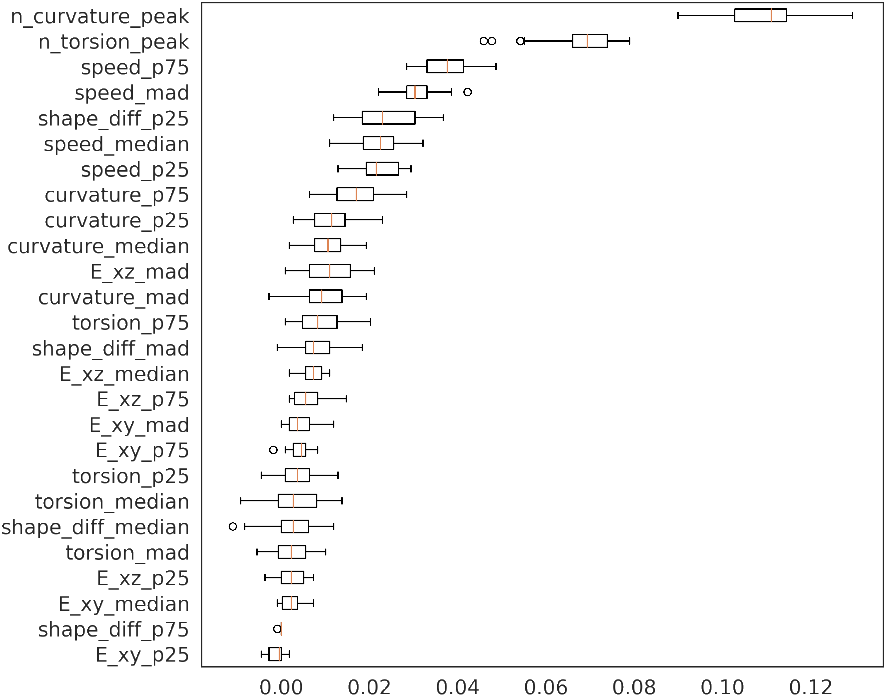
The permutation importance plot that shows the importance of each feature. We found that the movement features was dominantly selected as important features.

**Fig. 8.**
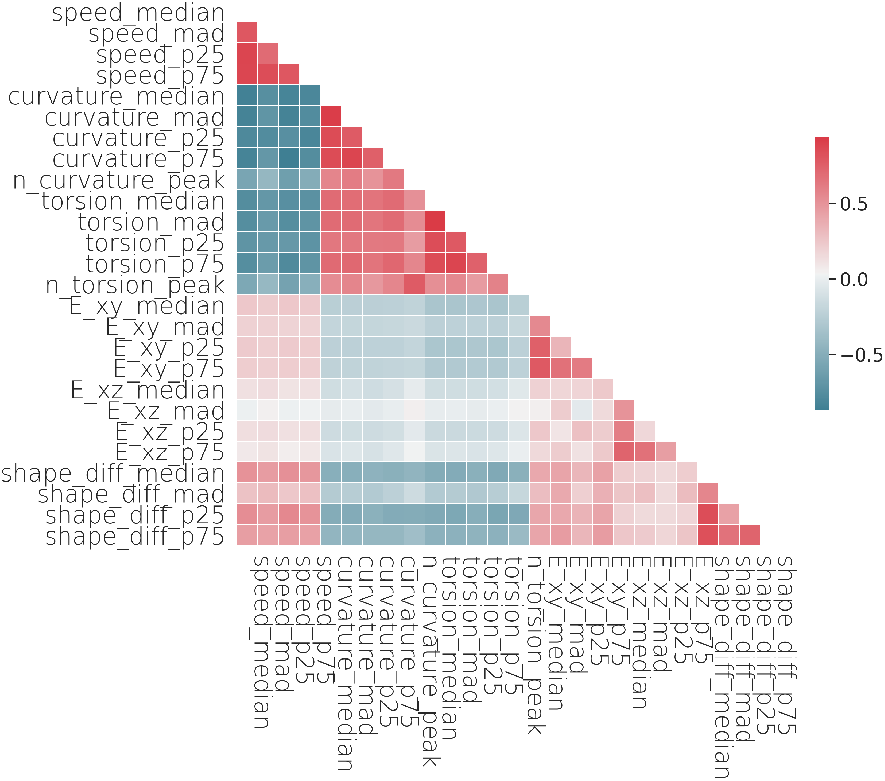
The correlation matrix of each features.

## CONCLUSION

In this paper, we have presented a novel application of parallel-transport moving frame to solve the shape alignment problem in 3D space and to integrate the analysis of shape and movement. We have shown that our approach can extract shape-movement features of the cell and their dynamic change. We found some interesting relationships between the movement and the cell shape such as the negative correlation between the cell speed and curvature/torsion, the fast cell tend to protrude the pseudopod to the movement direction, and positive correlation between shape change rate and the cell speed. The shape-movement features which are extracted using our approach also have the potential to be used for clustering or classification. It suggests that our approach can provide new insight into the mechanobiological process of the cell.

## Acknowledgements

We thank for Hironori Shigeta and Shigeto Seno of the Graduate School of Information Science and Technology, Osaka University, for the support in 3D mesh object creation.

## Contribution

Conceptualization: Y.D.H, R.Y., K.M.; Methodology: Y.D.H, R.Y.; Validation: Y.D.H, R.Y.; Formal analysis: Y.D.H; Investigation: Y.D.H, C.Y.C, Y.U.; Software: Y.D.H, C.Y.C; Data acquisition: Y.U., M.I.; Data curation: Y.U., M.I.; Writing - original draft: Y.D.H; Writing - review and editing: R.Y.; Visualization: Y.D.H; Supervision: R.Y.; Project administration: R.Y.; Funding acquisition: R.Y., M.I.

## Data availability

The source code and simulational dataset are available at https://github.com/yusri-dh/MovingFrame.jl.

## Supplementary

Movie 1

Movie 1. The simulational cell manually created using Blender.

Movie 2a

Movie 2a. The simulational cell shape before reorientation. The centers of mass for each timepoints are normalized to origin.

Movie 2b

Movie 2b. The simulational cell shape after reorientation. The centers of mass for each timepoints are normalized to origin.

Figure S2

The shape eccentricity calculated using SH coefficients.

## SUPPLEMENTARY MATERIAL

**Fig. S1.**
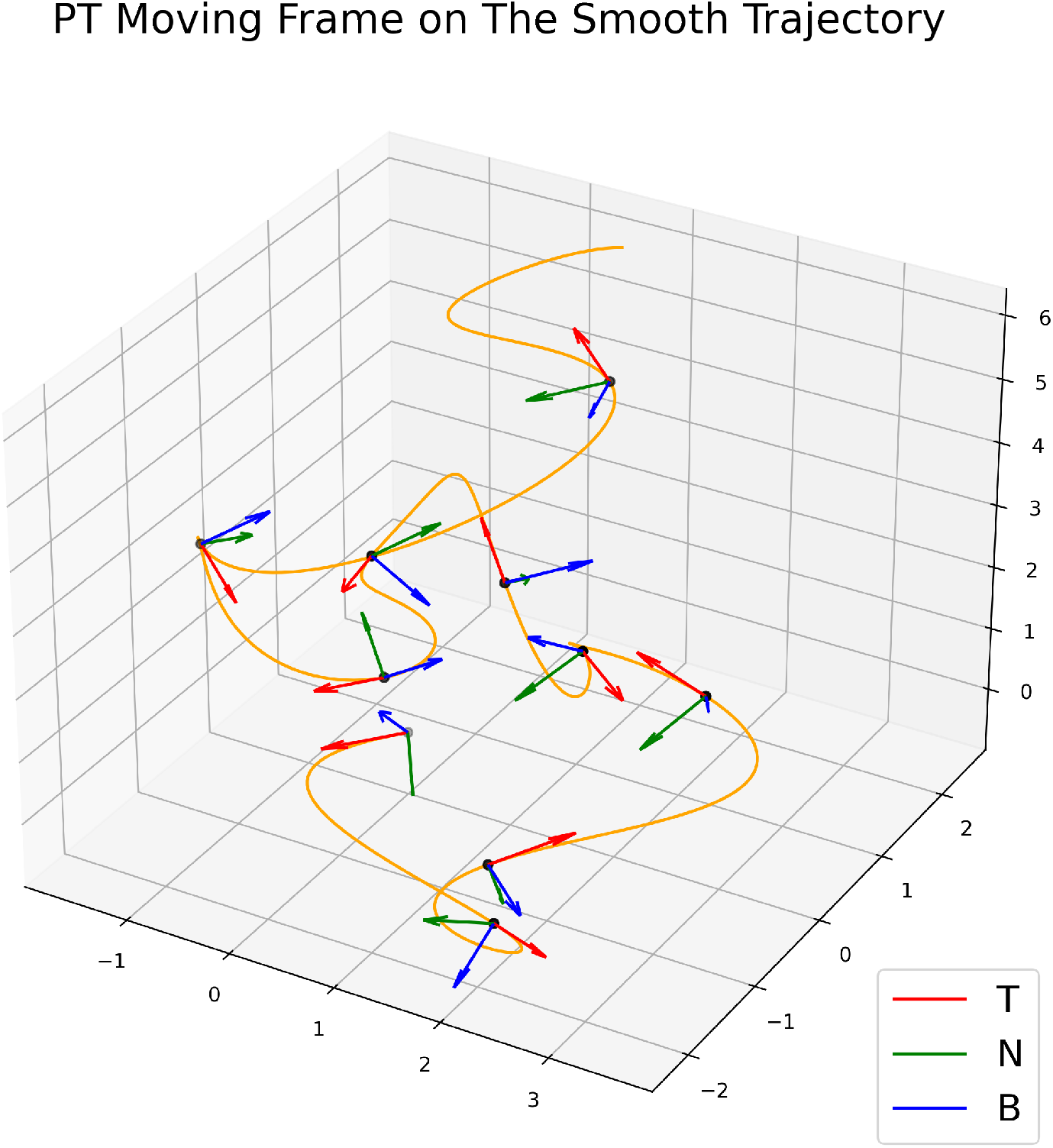
The construction of PT Moving Frame on each timepoints. The moving frames is moving with the curve and telling us the directions of the object movement.

**Fig. S2.**
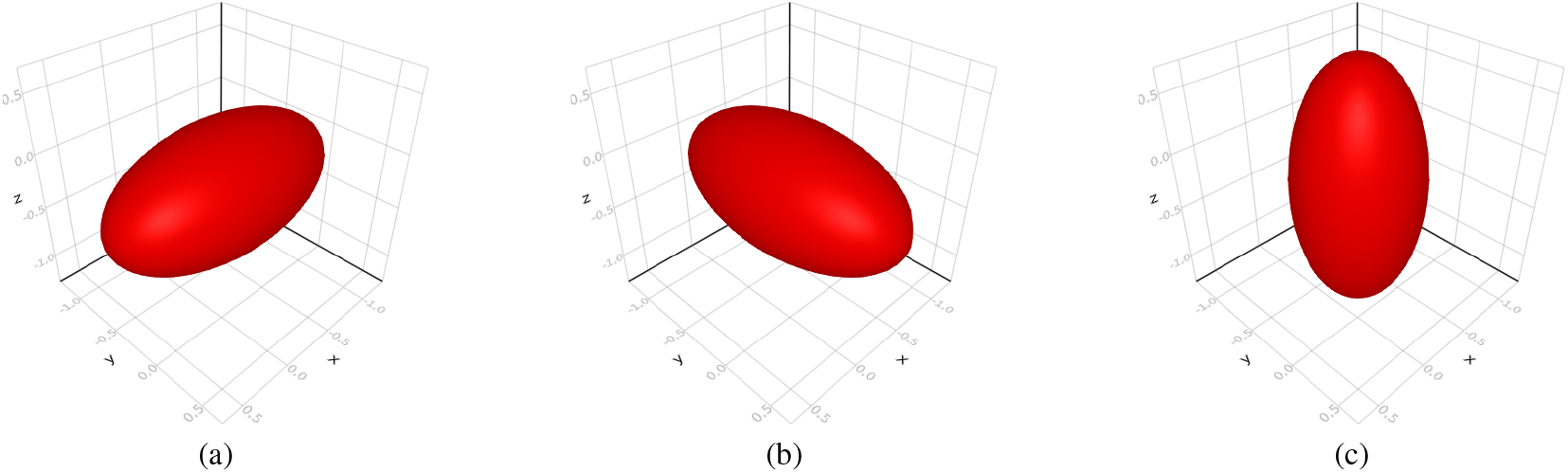
The shape eccentricity calculated using SH coefficients. For examples, figure (a) have eccentricity: *E_xy_* = 2, *E_xz_* = 2, *E_yz_* = 1, figure (b): *E_xy_* = 1/2, *E_xz_* = 1, *E_yz_* = 2, and figure (c): *E_xy_* = 1, *E_xz_* = 1/2, *E_yz_* = 1/2.

